# lncRNA *NORM* is essential for proper chromosome segregation through the Plk1-Bub1 and Nsun2 axis

**DOI:** 10.64898/2026.03.15.711899

**Authors:** Vikas Dongardive, Sonali Jathar, Juhi Srivastava, Vidisha Tripathi

## Abstract

The cell cycle comprises different phases and is a tightly regulated process at the molecular level. During the cell cycle, two key events occurred: DNA duplication during the S phase and chromosome segregation during mitosis. Accurate cell cycle progression, achieved through faithful chromosome segregation, is essential for maintaining cell fidelity. Long noncoding RNAs are a subclass of noncoding RNA that are longer than 200 bp and form RNA protein complexes (RNPs) to regulate various biological processes. Herein, we demonstrate that lncRNA *NORM* is involved in regulating the cell cycle by maintaining proper chromosome segregation. *NORM* exhibited G2 phase-specific expression, and the depletion of *NORM* resulted in a significant G2/M arrest. *NORM*-depleted cells failed to progress in mitosis and showed defects in chromosome segregation. We further demonstrated that *NORM* binds to proteins such as Plk1 and Nsun2. Depletion of *NORM* hindered the interaction between Plk1 and Bub1, resulting in reduced kinetochore localization of Plk1 during prometaphase. Our results also show that the depletion of *NORM* affects the binding of Nsun2 protein to CDK1 mRNA and, consequently, the stabilization of CDK1 at the protein level. Altogether, our results demonstrate that *NORM* regulates chromosome segregation by mediating the interaction between Plk1 and Bub1.

## Introduction

The mammalian cell cycle is a ubiquitous and tightly regulated process that comprises events orchestrated in an orderly manner to culminate in the formation of two daughter cells. The entry of a cell into different phases of the cell cycle is precisely controlled by the oscillating concentrations of different Cyclins and CDK complexes. In addition to Cyclins and CDKs, various cyclin kinase inhibitors (CKIs) and cell cycle regulatory proteins, such as RB, p53, and p21, regulate the cell cycle (1,2). Two key events that occur during the cell cycle are the duplication of genetic material in the S phase and the segregation of chromosomes in the M phase. Precise control of chromosome segregation during the M phase is crucial for error-free cell cycle progression, as any defect in chromosome segregation can lead to cellular abnormalities that ultimately contribute to the development of various cellular disorders, including cancer (3). One of the key events that is responsible for faithful chromosome segregation is the attachment of the kinetochore to microtubules (KT-MT) (4). A plethora of proteins are involved in mediating the proper attachment of KT-MT for the segregation of chromosomes. Any defect in chromosome segregation is detected by the surveillance mechanism known as the spindle assembly checkpoint (SAC), which is activated to correct the erroneous microtubule-to-kinetochore attachment (5). Budding uninhibited by benzimidazoles 1 (Bub1) plays a pivotal role in assembling the SAC complex by localizing at the kinetochore through the phosphorylation of Knl1 by Mps (6,7). Phosphorylated Knl1 then recruits the Bub3-Bub1 complex. Additionally, Mps1 phosphorylates Bub1 to mediate its interaction with the Mad1-Mad2 complex, thereby creating a SAC signaling platform (8). This platform then further recruits Bub3, BubR1, Cdc20, and Mad2. Bub1 also binds to another kinase, Polo-like kinase 1 (Plk1), and the interaction between Plk1 and Bub1 is essential for the kinetochore localization of Plk1. Any defects in Plk1 kinetochore localization cause chromosome mis-segregation, marked by chromosomes associated with spindle poles (9). The interaction between Bub1 and Plk1 primarily depends on Bub1’s phosphorylation at T609, mediated by the CDK1 protein (9). The Cyclin B and CDK1 complex drives the entry of a cell into mitosis and is crucial for guaranteeing accurate and faithful segregation of duplicated chromosomes. CDK1 is also regulated by various proteins, such as MYT1 and WEE1, which deactivate CDK1 through phosphorylation (10,11). Moreover, CDK1 is also regulated through mRNA stabilization by the protein binding to its mRNA. NOP2/Sun RNA methyltransferase family, member 2 (NSun2) (Misu) stabilizes CDK1 and enhances its translation by binding to its mRNA and methylating its 3’ untranslated region (UTR) (12). CDK1 binds to Cyclin B and becomes activated. Upon activation, it phosphorylates various substrates, including Bub1, which is crucial for mitosis and contributes to the faithful segregation of chromosomes. Multiple lines of evidence suggest that noncoding RNAs, including microRNAs (miRNAs) and long noncoding RNAs (lncRNAs), are also involved in regulating the cell cycle. lncRNAs are a class of noncoding RNAs that are longer than 200 bp and characterized by their short open reading frame. By forming RNP complexes, lncRNAs are known to control a variety of biological processes through Signal, decoy, Guide, or Scaffold mechanisms. In a recent study, it has been demonstrated that lncRNA MALAT1 is differentially regulated during the G1/S and M phases and is essential for the progression of the cell cycle from G1/S to S and from G2 to M, through the transcription and alternative splicing of the gene bMyb (13). Another study has demonstrated that the lncRNA EMS controls the cell cycle progression during the G1/S phase. EMS stabilizes E2F1 mRNA by binding to the RALLY protein in a manner that increases the expression of that gene (14). lncRNAs also interact with CDKs to regulate cell cycle progression. A recent study revealed that lncRNA HOXC-AS3 interacts with CDK2, thereby facilitating reduced binding of CDK2 to p21, which ultimately leads to increased CDK2 activity and promotes cell proliferation (15). Several other studies have demonstrated that many long noncoding RNAs (lncRNAs) play a role in cell cycle regulation. Among these lncRNAs, LINC00667 is an interesting lncRNA due to its recently studied role in multiple diseases, specifically in cancers and chronic kidney diseases. Recently, various studies have shown that LINC00667 is differentially expressed in various cancer cell lines and plays a crucial role in regulating cellular proliferation, invasion, migration, and apoptosis (16–19).Overexpression of LINC00667 in renal tubular epithelial cells results in chronic renal fibrosis by increasing cellular proliferation, migration, and apoptosis through the miR-19b-3p/LINC00667/CTGF axis(16). Upregulation of LINC00667 in clear renal carcinoma is associated with increased cellular proliferation, migration, and invasion, achieved through the targeting of miR-143-3p and a subsequent increase in ZEB1 expression (20). Furthermore, LINC00667 also plays an oncogenic role in various cancers, including Wilms’ tumor, through the miR-200b/c/429 and IKK-β axis (18). In non-small cell lung cancer, it plays its role via EIF4A3-stabilized VEGFA(17). In breast cancer, LINC00667 drives tumor growth and chemoresistance to docetaxel through the miR-200/Bcl2 axis (21). Additional evidence suggests that LINC006677 is upregulated in nasopharyngeal carcinoma, enhancing proliferation, migration, and the epithelial-to-mesenchymal transition by sponging miR-4319 and regulating the level of FOXQ1 (22). In esophageal cancer, LINC00667 is associated with promoting proliferation through the LINC00667/miR-200b-3p/SLC2A3 axis(23). In cholangiocarcinoma, LINC00667 regulates tumor growth through YY1-mediated upregulation (24). Together, these findings highlight LINC00667 as a crucial molecular regulator of proliferation in cancer and kidney disease, underscoring the need for further investigation to elucidate its mechanisms and therapeutic potential. Its consistent involvement in regulating cellular proliferation hints at a broader role in cell cycle control, an area that remains largely unexplored. Understanding this potential connection may uncover new therapeutic avenues for diseases where aberrant cell cycle regulation is a hallmark. Understanding the involvement of lncRNAs in cell cycle regulation will provide a detailed molecular mechanism and open up new opportunities for novel treatments of cellular disorders associated with cell cycle dysregulation.

In this study, we showed the involvement of LINC00667 in regulating the cell cycle. We demonstrated that long noncoding RNA LINC00667 is crucial for proper chromosomal segregation by modulating the interaction between Plk1 and Bub1. LINC00667 is a G2-phase enriched lncRNA and is essential for driving the G2/M progression of the cell. LINC00667 depletion causes G2/M arrest of cells marked with chromosome misalignment, which ultimately results in cell apoptosis. The interaction between Plk1 and Bub1 is hampered upon LINC00667 depletion. We have demonstrated the molecular relationship between LINC00667 and another protein, Nsun2. Additionally, we have demonstrated that LINC00667 depletion reduces the binding of Nsun2 to CDK1 mRNA, thereby decreasing the stability of the CDK1 protein. We observed that a reduced level of CDK1 protein results in a decrease in the phosphorylation of Bub1 at T609. Finally, Plk1 localization at the kinetochore is altered as a result of hindering the interaction between Plk1 and Bub1. Overall, our research suggests that the novel long non-coding RNA (lncRNA) LINC00667 plays a crucial role in regulating the cell cycle by ensuring normal chromosome segregation.

## Results

### *NORM* is upregulated during the G2 phase of the cell cycle and promotes G2/M transition

Tumour cells adapt pathways that bypass normal cell cycle checkpoints to enable unregulated cell division. Uncontrolled proliferation is often driven by signalling pathways that alter the expression of cell cycle regulators such as cyclins, cyclin-dependent kinases (CDKs), and CDK inhibitors. Therefore, oncogenic lncRNAs such as LINC00667, which promote proliferation, may exert their function at least in part by regulating cell cycle processes. Recent studies have demonstrated that one such long non-coding RNA (lncRNA), LINC00667, is differentially regulated in various cancers and plays a role in regulating proliferation. Uncontrolled cellular proliferation is one of the hallmarks of cancer, often resulting from defects in the cell cycle regulation program. The cell cycle is a more complex phenomenon that is coordinated through multiple layers of regulation, driven by proteins as well as a wide array of noncoding RNAs, including long noncoding RNAs (lncRNAs) and microRNAs (miRNAs). One of our comprehensive studies has identified a set of cell cycle-related long noncoding RNAs (lncRNAs), with most of these lncRNAs exhibiting cell cycle phase-specific expression patterns. LINC00667 is one such lncRNA that showed G2 phase-specific expression in synchronized HeLa cells. Based on its function, we named LINC00667 the <u>No</u>ncoding RNA <u>R</u>egulator of <u>M</u>itosis *(NORM). NORM* is approx. 4.0 nucleotides long noncoding transcript located on chromosome 18 (chr18:5,232,875-5,290,608 (GRCh38/hg38)), Histone tail modification map indicated a significant H3K4me3 and H3K27ac marks on the promoter of *NORM*, indicating that it is actively transcribed across different cell types **[FIG 1A].** The coding probability of the *NORM* transcript was analyzed using the CPAT tool, revealing that *NORM* does not code for any protein, unlike the well-characterized long noncoding RNA MALAT1 **[FIG. 1B].** To understand its cis-acting or trans-acting mode, *NORM* was further characterized for its subcellular localization using a subcellular fractionation method, followed by a quantitative real-time PCR analysis to assess the level of *NORM*. Quantitative real-time PCR data revealed that *NORM* is predominantly located in the nucleus, with some fraction also present in the cytoplasm **[FIG. 1C].** Quantitative real-time PCR (qRT-PCR) analysis revealed that the transcript levels of *NORM* remain low during the G1 and S phases of the cell cycle. However, there is a steady increase in transcript levels during the G2 phase, followed by a subsequent decrease in the M phase **[FIG. 1D].** Cell cycle phase-specific expression of lncRNAs is often associated with their role in that specific phase. An elevated level of *NORM* in the G2 phase suggests its potential role in regulating G2/M progression. To understand whether *NORM* is required for normal cell cycle progression and to examine the functional relevance of *NORM*, Antisense oligonucleotide (ASO)- based depletion of *NORM* was performed, followed by cell cycle analysis by using propidium iodide (PI) based fluorescence-activated cell sorting (FACS), which showed a significant G2/M arrest in asynchronously growing WI-38 cells **[FIG. 1E, F, and H]**. The effect of *NORM* depletion on cell cycle progression was further confirmed by performing BrdU-PI flow cytometry. We observed a significant increase in the G2/M population upon *NORM* depletion, confirming that *NORM* depletion leads to cell cycle arrest at the G2/M phase **[FIG. G]**. It was evident from the cell cycle analysis that the absence of *NORM* caused a significant G2/M arrest in the cell cycle, indicating the importance of *NORM* in driving cells through the G2/M phase. We further confirmed the role of *NORM* in driving G2/M progression by arresting the asynchronously growing HeLa cells in mitosis with Nocodazole treatment, followed by subsequent release in control and *NORM*-depleted conditions, and then analyzed by PI-FACS. PI-flow cytometry analysis indicated that in the control condition, cells were correctly arrested upon nocodazole treatment and subsequently progressed from the G2/M phase to the G1 phase upon release. Nevertheless, in the *NORM* depletion condition, cells failed to progress from the G2/M phase to the G1 phase even after 10 hours of release **[FIG. 1I & J].** Moreover, we observed that the level of CDK1 was reduced upon *NORM* depletion compared to the control, as determined by our immunoblot analysis **[Fig. 1K]**. This indicates the importance of *NORM* in driving the cells through the G2/M phase. As DNA content-based cell cycle analysis is incapable of distinguishing between the G2 and M-phase populations, we employed H3ser10 phosphorylation-based cell cycle analysis. A close examination of G2 and M phase populations revealed that *NORM* depletion results in a substantial decrease in the mitotic phase population and an increase in the G2 phase population, indicating a failure of the cells to progress from the G2 to the M phase **[FIG. 1 L&M]**. This was further confirmed by immunoblotting analysis, which checked the level of phospho-H3 at the ser10 residue. The level of phospho-H3 (ser10) was significantly reduced upon *NORM* depletion **[FIG. 1 N]**.

### Loss of *NORM* affects cellular proliferation, leading to an increase in apoptosis

Depletion of *NORM* resulted in a marked increase in the G2/M population, prompting us to investigate the effect of *NORM* on cellular proliferation. The effect on proliferation was evaluated by performing Ki-67 immunostaining. The results showed a considerable decrease in Ki-67-positive cell numbers in *NORM*-depleted conditions compared to control conditions **[FIG. 2A & B]**. *NORM* depletion resulted in a substantial reduction in cellular viability and proliferation, as well as a significant decrease in colony formation ability, as observed through the MTT assay and clonogenic assay, respectively, confirming the observations about proliferation **[FIG 2C, D, and E].** This data revealed that *NORM* depletion significantly reduced cell proliferation, suggesting that *NORM* depletion can lead to defects in cell cycle progression. The cell cycle analysis data showed an increase in the sub-G1 population upon *NORM* depletion, indicating a significant level of apoptosis in the absence of the lncRNA. We then determined the level of apoptosis in *NORM-*depleted cells by performing Annexin V-FITC and propidium iodide (PI) based flow cytometry. The flow cytometry data showed an increase in early and late apoptotic populations upon *NORM* depletion compared to those in control cells **[FIG. 2E & F].** Increased levels of cleaved PARP in *NORM-*depleted conditions further validated the result. Additionally, anti-apoptotic protein Bcl2 levels were decreased, while pro-apoptotic protein Bax levels were increased upon *NORM* depletion **[FIG. 2G].** The overall data suggest the importance of *NORM* in promoting cellular proliferation and inhibiting apoptosis.

### Loss of *NORM* results in changes in gene expression at the global transcriptomic and proteomic levels

To understand the molecular mechanism underlying the cell cycle defect upon *NORM* depletion, we analyzed gene expression changes by performing a global transcriptome analysis on control and *NORM*-depleted samples using the Illumina HiSeq 2500 platform. Global transcriptome profiling revealed that the loss of *NORM* results in the differential expression of 1,050 genes, wherein nearly 500 genes showed more than 2.5-fold up-regulation, while 550 genes showed more than 2.5-fold down-regulation **[FIG. 3A]**. ZNF48, SLC16A6, BRCA1, CHRND, PCNT, DCAKD, POT1, ZNF74, RNF14, and YY1AP1 were the top down-regulated genes, while SEC31A, ITGA2, FBXO6, DNM2, SMURF1, GUCD1, WDR4, POP1, PPP1R7, and PSMD1 were the top up-regulated genes **[Supplementary figure 1.1]**. Using qRT-PCR, we validated the expression of several differentially expressed genes (DEGs), which showed a similar trend **(Supplementary 1.2).** We next analysed the effect of *NORM* on cellular biological pathways by performing gene ontology analysis. Gene ontology revealed that many cell cycle-related biological processes and pathways were affected by *NORM* depletion. The top ten down-regulated biological processes and upregulated biological processes include cell cycle-related pathways such as regulation of mitotic cell cycle phase transition, regulation of cell cycle phase transition, positive regulation of cell cycle, G2/M transition of the mitotic cell cycle, and cell cycle G2/M phase transition **[FIG 3B &C]**. The down-regulation of G2/M transition-related genes, such as MELK, CEP70, CDK2, PPP2R2A, PCNT, and CDC25A, suggests a failure in the G2/M transition in the absence of *NORM*. To gain further insight, we performed a proteomics analysis in *NORM-*depleted and control conditions to check the effect of *NORM* depletion on different proteins and biological processes. *NORM* depletion resulted in > 2-fold down-regulation of 146 proteins and > 2-fold up-regulation of 679 proteins **[FIG 3D].** Many cell cycle-related proteins showed differential expressions. Proteins known for their role in cell cycle regulation, such as TGM2, PSMD, CLTC, PSMD2, BUB3, and ANXA4, were significantly upregulated. In contrast, proteins such as PDGFRB, PTMS, COL1A1, CD2AP, LAMB2, YWHAB, CDK1, MCM3, and YWHAG were significantly reduced upon *NORM* depletion **[Fig. 3E].** To further understand this, we performed a Gene Ontology (GO) analysis of proteomics data using the DAVID tool. The gene ontology analysis of the proteomics data revealed that biological processes, such as “negative regulation of ubiquitin-protein ligase activity involved in mitotic cell cycle, anaphase-promoting complex-dependent catabolic process, mitotic nuclear envelope disassembly, mitotic spindle assembly, Cell cycle, and G2/M transition of the mitotic cell cycle,” were affected **[Fig. 3F].** The Gene Ontology data suggests that *NORM* is involved in regulating mitosis-related pathways. Global transcriptome and proteomics profiling data indicate an overlap between RNA-seq and proteomics profiles. Altogether, our high-throughput data suggests the importance of *NORM* in regulating the cell cycle by associating with mitosis-related pathways.

### *NORM* interacts with cell cycle regulatory proteins for chromosome segregation and cell cycle regulation

We next sought to investigate the mechanism of action of *NORM* in regulating the cell cycle. We incubated the *in vitro* transcribed full-length biotinylated *NORM* with cell lysate, followed by a pulldown of the *NORM*-protein complex using streptavidin beads **[FIG. 4A]**. *NORM* showed interaction with numerous proteins, including cell cycle regulatory proteins CALM2, CDC5L, SPOUT1, KHDRBS1, Nsun2, and Plk1, out of which polo-like kinase 1 (Plk1) seemed to be interesting, as Plk1 is required for entry of cells into mitosis. Our previous results showed a defect in the G2/M transition under NORM depletion conditions [FIG. 4B & **C]**. The *NORM*-Plk1 interaction was confirmed by performing RNA immunoprecipitation, which showed significant enrichment (∼3-fold) of *NORM* in the Plk1 immunoprecipitated **[FIG. 4D]**. Interestingly, we also observed a marginal decline in Plk1 protein levels upon *NORM* depletion **[FIG. 4E]**. Plk1 is one of the major kinases in mitosis, phosphorylating a multitude of substrates at various subcellular locations to regulate mitotic events. Plk1 is essential for the proper assembly of spindles, centrosome maturation, and the attachment of microtubules to kinetochores. We then investigate the role of *NORM* on Plk1’s localization at different subcellular locations. We observed differential localization of Plk1 at various subcellular locations, with some defects, in the *NORM*-depleted condition **[FIG. 4F].** To understand the role of *NORM* on Plk1 localization at kinetochores and centrosomes, we examined the level of Plk1 at kinetochores in control versus *NORM* depletion conditions using immunofluorescence. We observed a significantly reduced intensity of Plk1 at kinetochores upon *NORM* depletion compared to control cells, suggesting that *NORM* depletion affected Plk1 localization at kinetochores **[FIG. 4 G&H].** We next sought to determine the effect of *NORM* depletion on chromosome segregation. We observed that depletion of *NORM* leads to defective chromosome segregation, marked by the misalignment of chromosomes during mitosis, and a significantly higher number of cells showed lagging chromosomes at the spindle pole **[Fig. 4 I & J]**.

### *NORM* depletion hampered the interaction between PLK1 and Bub1

Next, to elucidate the mechanism of reduced localization of Plk1 at kinetochores, we examined the effect of *NORM* depletion on the interaction of key regulatory proteins involved in mitosis. With the help of string analysis, we found that many cell cycle regulatory proteins, such as cyclin B, CENP-E, and Bub1, are known to interact with Plk1 **[FIG. 5A]**. The interaction between Bub1 and Plk1 is crucial for the localization of Plk1 at kinetochores. Plk1 interacts with Bub1 through its polo-box domain (PBD) and phosphorylates Bub1. First, we checked if there was any effect of *NORM* depletion on the level of Bub1. We checked the kinetochore localization of Bub1 in control and *NORM*-depleted cells. We observed a marginal change in Bub1 intensity at the kinetochore upon *NORM* depletion **[FIG. 5B & C]**, and no significant change was observed at the protein level of Bub1 upon *NORM* depletion **[FIG. 5D]**. We then checked the effect of *NORM* on the interaction between Plk1 and Bub1. Interestingly, when we immunoprecipitated Plk1, it was able to co-immunoprecipitate Bub1 along with it; however, upon *NORM* depletion, the interaction between Plk1 and Bub1 was hampered, as Plk1 failed to co-immunoprecipitate Bub1 **[FIG. 5E]**. The interaction was validated by performing reverse Co-immunoprecipitation in which an antibody against Bub1 was used to immunoprecipitate Bub1. Reverse Co-IP also showed that upon *NORM* depletion, the interaction between Plk1 and Bub1 was hampered **[FIG. 5F]**. It is known that CDK1 phosphorylates Bub1 at the T609 residue, and this phosphorylation is essential for the interaction between PLK1 and Bub1 (25). A mitosis-specific anti-MPM2 antibody can be used to detect Bub1 phosphorylated at the T609 residue, and phosphorylation of Bub1 at T609 is crucial for the Plk1-Bub1 interaction and the localization of Plk1 at the kinetochore (26). Considering the importance of Bub1 T609 in Plk1 interaction, we checked the effect of *NORM* depletion on Bub1 phosphorylation. We immunoprecipitated Bub1 using an anti-Bub1 antibody and assessed the phosphorylation status of Bub1 at T609 using an anti-MPM2 antibody. Interestingly, we observed that *NORM* depletion led to a significant reduction in Bub1 phosphorylation at the T609 residue **[FIG. 5G & H]**, indicating that *NORM* is involved in mediating Bub1 phosphorylation. The depletion of *NORM* resulted in a decrease in Bub1 phosphorylation, ultimately hindering Bub1’s interaction with Plk1. Altogether, these results suggest that the *NORM* is involved in mediating the interaction between Plk1 and Bub1 through the phosphorylation of Bub1 at T609.

### *NORM* mediates the binding of Nsun2 on CDK1 mRNA to stabilize CDK1 protein

We investigated Bub1’s upstream targets to gain a deeper mechanistic understanding of how *NORM* regulates the phosphorylation of Bub1. It is known that Bub1 is phosphorylated at T609 by cyclin-dependent kinase 1 (CDK1). Therefore, we explore any potential association between *NORM* and CDK1. Our RNA pulldown data showed that *NORM* lncRNA also interacts with the tRNA methyltransferase Nsun2 (Misu). We validated this interaction by performing an RNA pulldown followed by western blotting to verify the presence of Nsun2 **[Fig. 6A]**. Nsun2 binds to CDK1 mRNAs. It stabilizes them at the protein level by methylating them. Next, we examined the effect of *NORM* depletion on Nsun2 at the protein level. We found no apparent change in Nsun2 at the protein level, indicating that *NORM* does not directly control Nsun2 expression **[Fig. 6B].** Next, we examined the effect of *NORM* depletion on Nsun2 binding to CDK1 mRNA with RNA-immunoprecipitation **[Fig. 6D]**. Interestingly, our RNA-IP with anti-Nsun2 antibody showed that upon depletion of NORM, CDK1 enrichment on Nsun2 was considerably lower than in control cells **[Fig. 6D].** These results suggested that *NORM* depletion inhibits Nsun2’s propensity to bind to CDK1 mRNA. Next, we checked the effect of reduced binding on the CDK1 protein level. Immunoblotting data suggested that the CDK1 was reduced at the protein level upon NORM depletion [Fig. 6E], suggesting that the reduced level of Nsun2 binding of CDK1 mRNA was hampered by *NORM* depletion, and hence it resulted in reduced stability of CDK1 protein. This reduced level of CDK1 resulted in decreased phosphorylation of Bub1 at T609, ultimately hindering the interaction between Plk1 and Bub1. Overall, the evidence suggests the necessity of *NORM* for chromosomal segregation and normal cell cycle progression by mediating Nsun2’s binding to CDK1 mRNA and modulating the interaction between Bub1 and Plk1.

## Discussion

In this study, we have demonstrated that *NORM*, a long noncoding RNA, plays a crucial role in regulating cell cycle progression from the G2 to M phase. We observed that *NORM* shows an elevated level of expression in the G2 phase. Multiple studies have demonstrated that lncRNAs exhibit tissue- and cell-stage-specific expression. This tissue- and cell-specific expression is often associated with the function of lncRNAs in that cell or tissue. The Kannanganattu V Prasanth group has demonstrated that more than 2,000 long noncoding RNAs (lncRNAs) exhibit periodic expression during the cell cycle. (27). An elevated level of lncRNA is found to be associated with cell stage-specific regulation of the cell cycle phases in a cell stage-specific manner. Therefore, this periodic expression of *NORM* may play a crucial role in regulating the cell cycle. Indeed, we found that when we depleted *NORM* cells, they were arrested at the G2/M phase and were unable to move to the G1 phase upon release. The arrest of cells may be due to the defective assembly of proteins required for mitotic progression. Bida *et al.* also demonstrated that a mitosis-associated long noncoding RNA (lncRNA), MA-linc1, is essential for mitosis. In the absence of lncRNA-MA, cells are unable to progress from mitosis to the G1 phase of the cell cycle.

We evaluated the role of *NORM* in regulating cellular proliferation and found that *NORM* plays a crucial part in this process, with depletion of *NORM* resulting in reduced cell proliferation. The G2 phase is a critical phase for cell proliferation, as it determines the cell’s fate, whether it will progress into the M phase and divide into two daughter cells. Other evidence also suggests that lncRNA depletion causes cell cycle arrest, leading to a decrease in cellular proliferation.(28). Reduced cellular proliferation was accompanied by an increase in the rate of apoptosis and an increase in the level of DNA damage. When a cell is arrested at any checkpoint for an extended period, the apoptotic cascade is activated. We also observed that cells were arrested for a longer time in the G2/M phase. This arrest may have triggered the apoptotic cascade, leading to an increased rate of apoptosis.

At the molecular level, a cell’s entry into each phase is regulated by various types of cyclins, cyclin-dependent kinases (CDKs), cyclin kinase inhibitors, and specific transcription factors. We observe a significant change in the protein level of CDK1 upon *NORM* depletion. CDK1 is a key regulator of the G2 and M phases of the cell cycle. It, along with cyclin A and Cyclin B, regulates cell entry from the G2 phase to the M phase (29–32). A decrease in the level of CDK1 is one possible reason cells are arrested at the G2/M boundary, and this defect also causes defects in mitotic cells.

Through RNA sequencing analysis and proteomics analysis, we found that pathways such as the Regulation of mitotic cell cycle phase transition and the G2/M transition of the mitotic cell cycle were affected. This suggests that *NORM* plays a crucial role in cell cycle progression. DE mRNAs, such as BRCA1, FBXO6, and PCNT, which are involved in cell cycle regulation, were significantly affected. BRCA1 plays a crucial role in regulating the cell cycle(33,34). PCNT is a significant protein that helps in assembling the PCM complex for centrosome maturation(35–37). Therefore, alterations in various genes related to cell cycle progression could have resulted in changes to biological processes associated with the cell cycle. Proteomics analysis also demonstrated that many proteins and the pathways related to cell proliferation and cell cycle were affected.

In this study, we observed a significant decrease in Plk1 localization at the kinetochore. At kinetochore Plk1 interacts with Bub1, BubR1 and MAD2 proteins(38–41). The interaction of Plk1 with these proteins is essential for the nucleation of PCM components, including PCNT (36,42). The interaction of Plk1 at the kinetochore facilitates the attachment of the kinetochore to the microtubules (MT). Lera et al. observed a similar result, showing that Plk1 inhibition can cause chromosome missegregation, characterized by lagging chromosomes (43). Therefore, the reduced Plk1 localization at the kinetochore may have impacted the target proteins CENP-U, BubR1, PICH, CLIP-170, CLASP2, Survivin, USP16, and RSF1, for example, including Sgt1 phosphorylation, which could result in improper attachment of microtubules to the kinetochore.(44). PLK1 phosphorylates CENP-Q by binding to CENP-U, and the absence of phosphorylation can lead to chromosome missegregation(45).

The interaction between Bub1 and Plk1 is crucial for the proper segregation of chromosomes during cell division. In our study, we observed that *NORM* depletion hampers the interaction between these two proteins. It has been reported in studies that Bub1 becomes phosphorylated at its T609 residue by CDK1, a crucial event for the binding of Bub1 to the PBD domain of Plk1. So, the reduced interaction between Bub1 and Plk1 is a consequence of the reduced CDK1-mediated phosphorylation on Bub1 T609. Plk1 also interacts with CENP-U, along with Bub1, at the kinetochore for chromosome segregation; however, this study has not explored the effect of the interaction between CENP-U and Plk1. In our study, we observed that in the absence of *NORM*, the binding of Nsun2 on CDK1 mRNA was significantly reduced. Other studies have also demonstrated the importance of Nsun2 binding to CDK1 mRNA and its significance in the translation of CDK1 mRNA by mediating methylation. Although our data suggested a direct interaction between the Nsun2 protein and *NORM*, we did not observe a direct interaction between NORM and CDK1 mRNA, indicating that *NORM* lncRNA and CDK1 mRNA do not form an RNA-RNA interaction complex. This suggests possibilities other than a direct RNA-RNA interaction. One of the limitations of our study is that we have not explored in detail how exactly the *NORM* mediates the interaction between Nsun2 and CDK1 mRNA, nor whether there is any change in the methylation of CDK1 mRNA. However, we speculate that *NORM* acts as a Guide lncRNA for Nsun2 protein to mediate the methylation of CDK1 mRNA, thereby increasing its translation and subsequently phosphorylating Bub1 at T609. Various studies have shown that lncRNAs can exert their functions by acting as a guide molecule for the protein (46).

## MATERIALS AND METHODS

### Cell culture

HeLa cells and Wi-38 cells were cultured in DMEM and MEM, respectively, supplemented with high glucose and 10% fetal bovine serum (FBS), as well as a penicillin-streptomycin antibiotics solution (Gibco). Cells were maintained at 37°C in a humidified atmosphere of 5% CO2.

### Antisense oligonucleotide treatment

Transient transfection of cells was performed using. For transfection, LNA GapmeRs Phosphorothioate internucleosidic linkage-modified antisense oligonucleotides were diluted in Opti-MEM. *NORM* Gapmers were used at a final concentration of 100 nM, twice (48 hours apart) within a 24-hour gap, to deplete Human *NORM*.

### Flow cytometry

PI-Flow cytometry was used to analyze the distribution of cell cycle phases. Cells were harvested using a scraper and then collected by centrifugation. Cells were washed with 1X PBS (phosphate-buffered saline) and fixed in ethanol for at least 4 to 5 hours or overnight at 4 °C. Cells were washed two times with 1XPBS and then incubated in 500ul of 1X PBS containing 0.1% Triton X-100, 20mg/ml propidium iodide, and 0.01mg/ml RNase A for 30 min at 37°C. It was followed by cell cycle analysis using a FACSCalibur (BD Bioscience), and the data were analyzed using CellQuest Pro.

For BrdU-propidium iodide (PI), flow cytometry was performed as described earlier (Tripathi et al., 2013). Briefly, S-phase cells were labeled with BrdU at a final concentration of 50 µM and incubated for 1 hour at 37°C. Cells were further resuspended in chilled 0.9% NaCl and then fixed in an equal volume of ethanol, followed by incubation at −20°C for 1 hour. Furthermore, cells were resuspended and incubated in a 2N HCl + 0.5% Triton X-100 solution for 30 minutes at room temperature, followed by washing in 0.1 M Sodium tetraborate solution. Finally, the cells were resuspended in PBS/0.5% Tween-20 + 1% BSA and incubated with the anti-BrdU antibody for 1 hour at RT. The cells were incubated in PBS + PI/Triton X-100 at 37°C for 15 min. Cells were analyzed using a FACScalibur flow cytometer (BD Bioscience), and the data were analyzed with CellQuest Pro.

For mitotic cell counting, total cells were collected by trypsinization. It was followed by fixing the cells in chilled 70% ethanol. Cells were permeabilized by using 0.25% Triton X-100. Cells were incubated with H3ser10 antibody (Abcam, ab14955) for two hours at room temperature. It was followed by brief washing and blocking of the cells with 1% BSA. Further, cells were incubated with Alexa Fluor 488 secondary antibody for 30 minutes at room temperature. Moreover, finally, cells were incubated with Propidium iodide (PI) containing RNase for 30 minutes. Cells were analyzed using a FACScalibur flow cytometer (BD Bioscience), and the data were analyzed with CellQuest Pro.

To detect apoptosis, dual staining with Annexin V and propidium iodide (PI) was performed according to the manufacturer’s instructions (Invitrogen, cat. no R37174). Briefly, Wi-38 cells were trypsinized, collected, and washed with ice-cold PBS. Cells were resuspended in 500 μL of 1x binding buffer. A drop of Annexin-V FITC was added, and cells were incubated for 15 minutes at 25°C in the dark. 5 μL propidium iodide was added before acquiring the cells. Cells were acquired using a BD FACScanto, and data were analyzed with CellQuest Pro.

### Total RNA extraction and quantitative real-time PCR (qRT-PCR)

Total RNA was isolated from cells using Trizol reagent (Invitrogen) according to the manufacturer’s instructions, and the concentration was estimated using a UV spectrophotometer. Genomic DNA was digested by DNase I (Sigma) treatment. Total RNA was subjected to first-strand cDNA synthesis using Multiscribe Reverse Transcriptase and Random Hexamers (Invitrogen). Transcript levels were quantified by quantitative real-time PCR (qRT-PCR) using gene-specific primers.

### RNA-seq analysis

Whole-transcriptome analysis of control and *NORM*-depleted samples was performed on the Illumina HiSeq 2500 platform to achieve high transcriptome coverage (50-100 million reads per sample). Briefly, Total RNA was isolated by the RNeasy kit (Qiagen). RNA-seq libraries were constructed using total RNA isolated from untreated and *NORM*-depleted samples with the RNeasy kit (Qiagen). Samples were indexed with adaptors and submitted to paired-end sequencing. RNA-seq reads were aligned with Tophat2.0.11 to the human genome. A comparison of expression levels was calculated by using fragments per kilobase of transcripts per million fragments mapped (FPKM). Gene ontology analysis was performed by using the Database for Annotation, Visualization, and Integrated Discovery (DAVID) tool (https://david.ncifcrf.gov).

### Quantitative proteomics by label-free quantitation mass spectrometry

Human lung fibroblast primary cells (WI38) were seeded in 10 cm cell culture plates for both the scrambled and NORM-depleted conditions. Cells were collected and pelleted in 1X PBS. 1X PBS was removed entirely, and the cell pellet was snap-frozen by liquid nitrogen for 20–30 seconds and stored at -80 °C. Urea lysis buffer was used for cell lysis, and the cells were sonicated to ensure complete lysis. Acetone precipitation was performed by incubating overnight at -20 °C with acetone. Urea rehydration buffer was used for the elution of proteins, followed by the removal of salts. An Amicon ultra-0.5 centrifugal filter unit with a 3 kDa cut-off (UFC500394) was used to remove salt and concentrate the sample. Protein was estimated using Pierce reagent, and bovine serum albumin (BSA) was used for plotting the standard curve. An equal amount of protein (50 µg) was digested using the TFE-In solution digestion protocol. Desalting and concentrating peptides or small proteins were performed using a ZipTip with 0.6 µL C18 resin (ZTC18S096). Mass spectrometry sample preparation was performed by adding 0.1% formic acid to dried protein samples. Samples were acquired by injecting a 2 µL sample into the Thermo Scientific Orbitrap Fusion Tribrid LC-MS/MS system. Raw files were transferred to the Proteome Discoverer software for analysis of the sample.

### Cell proliferation assay

Cell viability was assessed using the 3-(4,5-dimethylthiazol-2-yl)-2,5-diphenyltetrazolium bromide (MTT) assay. Briefly, after transfection with *NORM* ASO or scrambled oligos for 48 hours, HeLa cells were trypsinized, and a suspension of 5000 cells was seeded into a 96-well plate. After 24 hours, 20 μL of MTT (5 mg/mL) was added, and the cells were incubated for an additional 4 hours. Then, 150 μL of isopropanol was added to dissolve the crystals. The proliferation ability of cells was assessed at 0 hours, 24 hours, 48 hours, 72 hours, and 96 hours by incubating the cells at 37°C. Absorbance at 570 nm was determined spectrophotometrically by using a microplate reader. Colony-forming ability was checked by the colony formation assay. Briefly, 1,000 cells were seeded in a six-well plate and maintained for 2 weeks, during which the media were replaced every 3 days. Colonies were fixed by PFA and stained by using crystal violet in PBS for 30 minutes. The colony formation ability was assessed by counting stained colonies.

### Immunoblotting

Cells were washed twice with PBS and then collected by scraping. A lysis buffer containing protease and phosphatase inhibitor was used for extraction. Loading dye was added directly to the samples, and the samples were denatured by boiling for 5 min at 95°C. The total cell lysate was subjected to SDS-polyacrylamide gel electrophoresis (SDS-PAGE), and proteins were transferred to the PVDF membrane. The membrane was incubated with primary antibodies overnight at 4°C. An appropriate secondary antibody was incubated for 1 hour at room temperature. The antibodies used for immunoblotting were anti-CDK1(Cdc2) (1:1000; Santa Cruz SC-54), anti-PARP (1:1000; Cell signaling technology, 9542), anti-Bax (1:500; Santa Cruz sc493), anti-Bcl2 (1:500; Santa Cruz sc492), anti-Plk1 (1:1000 Abcam ab17056), anti-Cyclin A (1:1000; Santa Cruz sc271682), anti-Cyclin B(1:1000; Santa Cruz sc7393), anti-Cyclin E (1:1000; Santa Cruz sc377100), anti-CDK4(1:1000; Santa Cruz sc23896), anti-Bub1(1:100 ab54893), H3ser10 antibody (1:1000; ab14955;), anti-α-Tubulin (1:5000; Sigma T6199), Anti-mouse IgG, HRP-linked (1:5000; cell signalling technology; 7076), Anti-rabbit IgG, HRP-linked (1:5000; cell signalling technology; 7074).

### Immunostaining

Immunofluorescence was performed as described earlier (Tripathi et al., 2013). Briefly, Wi-38 cells, grown on coverslips, were then fixed in 2% PFA for 20 minutes at room temperature (RT) and permeabilized with 0.1% Triton X-100 for 10 minutes on ice. Cells were blocked by incubating with 5% fetal bovine serum (FBS) at room temperature (RT) for 1 hour, followed by incubation with the primary antibody for 2 hours at RT. Incubation with fluorescence-tagged secondary antibody was performed for 1 hour at room temperature. Finally, cells were stained with Hoechst and mounted in mounting media. The antibodies were used are Ki-67 (1:500; Abcam; ab16667), anti-Plk1 (1:100 Abcam; ab17056), anti-α-Tubulin (1:1000; Sigma T6199), anti-Bub1(1:100; Abcam; ab195268), anti-Pericentrin (1:200; Novus; NB 100-61071) CREST (1:500; antibodiesinc; 115-234), Alexa flour 488 donkey anti-rabbit IgG (H+L) (1:1000; Invitrogen; A21206), Alexa flour 594 donkey anti-mouse IgG (H+L) (1:1000; Invitrogen; A21203) images were acquired on Zeiss microscope.

### RNA pulldown and Mass spectrometry analysis

*NORM* was cloned into a pGEMT vector. The plasmid was linearized by using the NcoI restriction enzyme. Purified plasmids were then used for in vitro transcription with a Biotin RNA labeling Mix (Roche). Biotin labeling was done by using T7 polymerase at 37°C for 6 hours. In vitro transcribed RNA was then purified by passing it through a MicroSpin column (Illustra MicroSpin G-25 columns, 27-5325-01, Merck). Fresh lysate was prepared by using Protein extraction buffer (20 mM Tris HCl, pH 7.5, 100 mM KCl,5 mM MgCl2,0.5% NP40) with protease, phosphatase cocktail inhibitors, and RNase inhibitors. The lysate was precleared using magnetic streptavidin Dynabeads (M-280 streptavidin, Cat. No. 11205D, Invitrogen). Biotin-labeled RNA and lysate were incubated on a rotator for 2 hours. It was followed by the addition of magnetic streptavidin Dynabeads to a mixture of RNA-lysate. It was further incubated for 2 hours. Next, protein samples were separated by SDS-PAGE, and bands were visualized by silver staining. For mass spectrometry analysis, the immunoprecipitated protein was washed with 100 mM ammonium bicarbonate. Proteins were prepared for mass spectrometry by digesting with RapiGest at 60°C for 30 minutes and then analysed by mass spectrometry.

### RNA Immunoprecipitation

Cells were washed twice with ice-cold phosphate-buffered saline (PBS). Cells were irradiated with 300mJ/cm^2^ in a UV crosslinker. The cell lysate was prepared by using RSB100 lysis buffer (100 mM Tris-Cl, pH 7.4, 100 mM NaCl, 2.5 mM MgCl_2_, 0.5% NP40, 0.5% Triton X-100) by incubating on ice for 15 minutes. The lysate was centrifuged, and the supernatant was collected. 0.1M DTT, RNase Inhibitor, and DNase I were added to the supernatant. The lysate was precleared by incubating Protein G agarose beads (Pierce, Cat. No. 20398) at 4°C for 1 hour, rotating on a rotor shaker. Immunoprecipitation was performed by incubating an equal amount of lysate with an anti-Plk1 antibody (1:100; Abcam, ab17056), an anti-Nsun2 (1:100; Abcam, ab259941), and a mouse IgG antibody (Negative control) overnight on a rotator shaker at 4 °C. Next, protein G agarose beads were added and incubated further for 2 hours. Co-immunoprecipitated RNA was isolated by resuspending the beads in 1 mL of Trizol (Invitrogen). RNA was isolated according to the manufacturer’s instructions. cDNA was prepared using 5X PrimeScript RT Master Mix (Takara, cat. no RR036A-1).

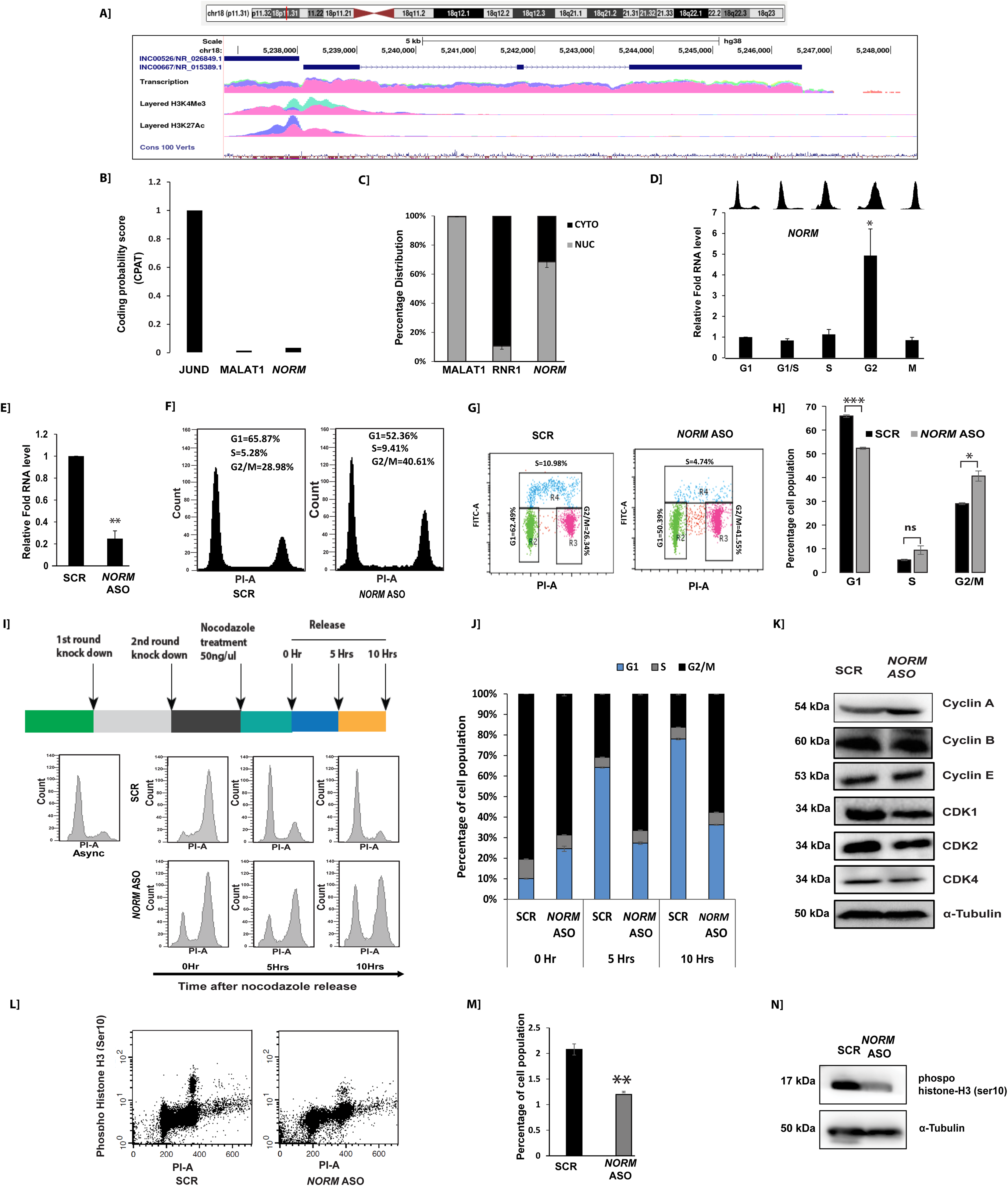

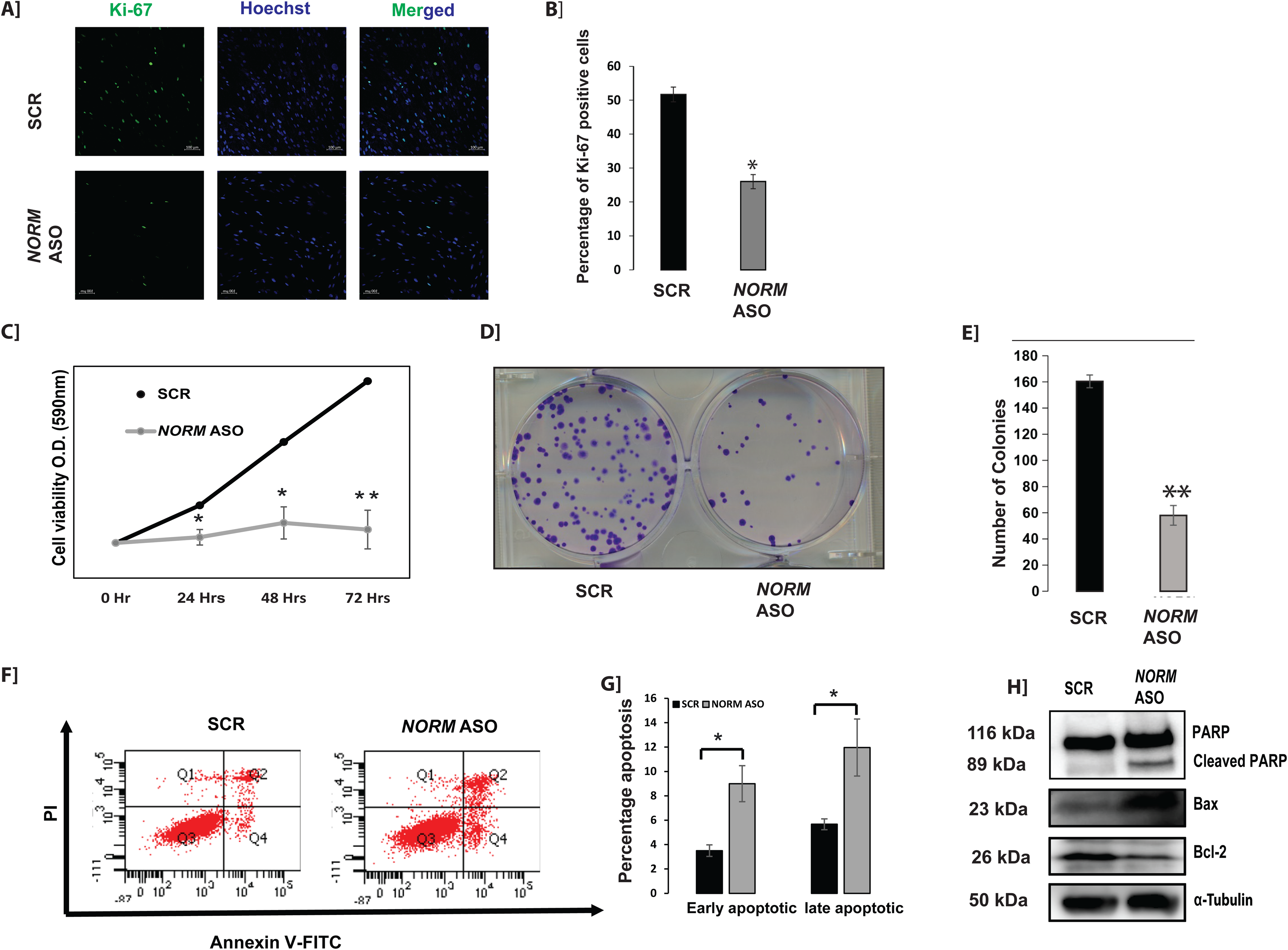

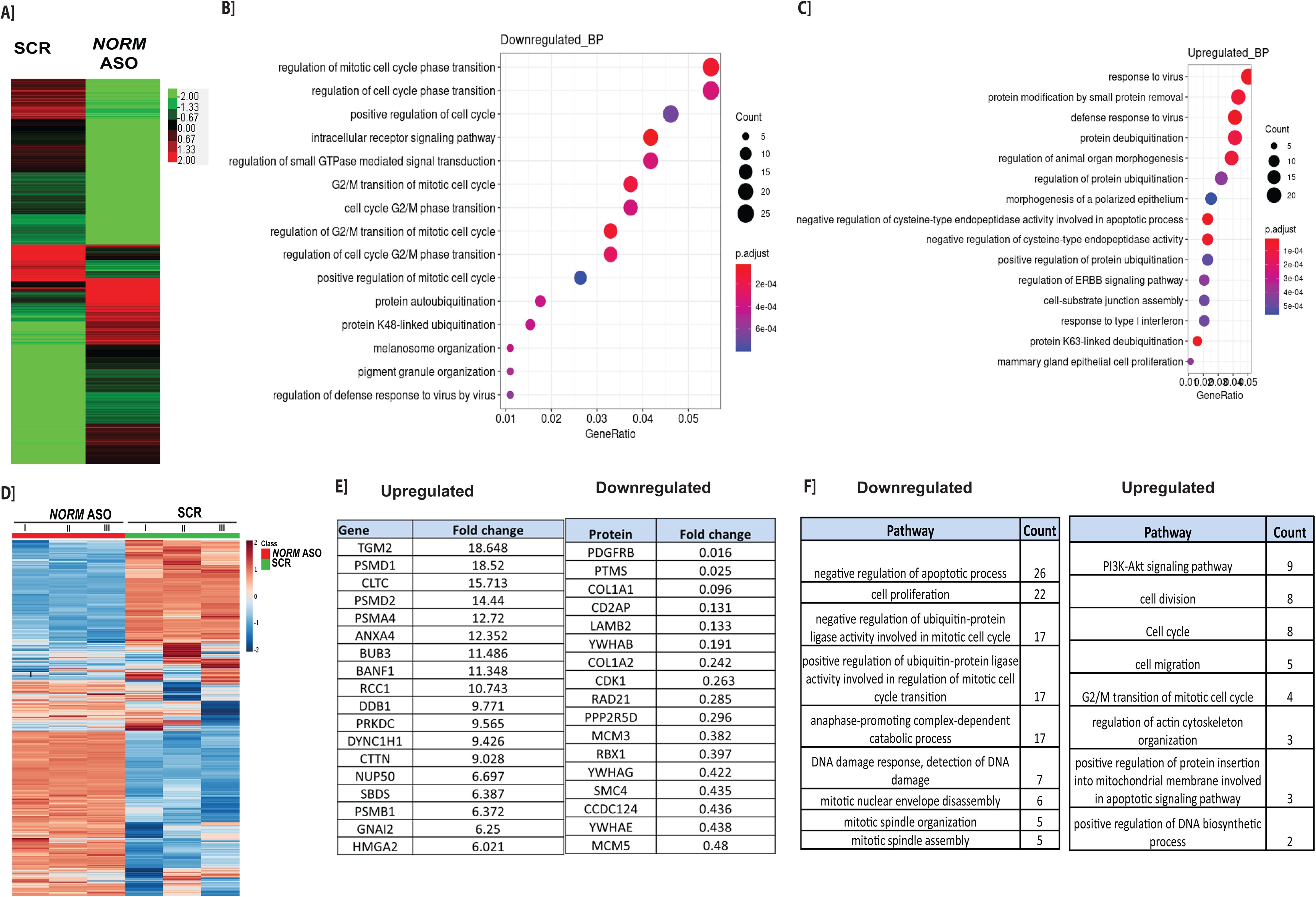

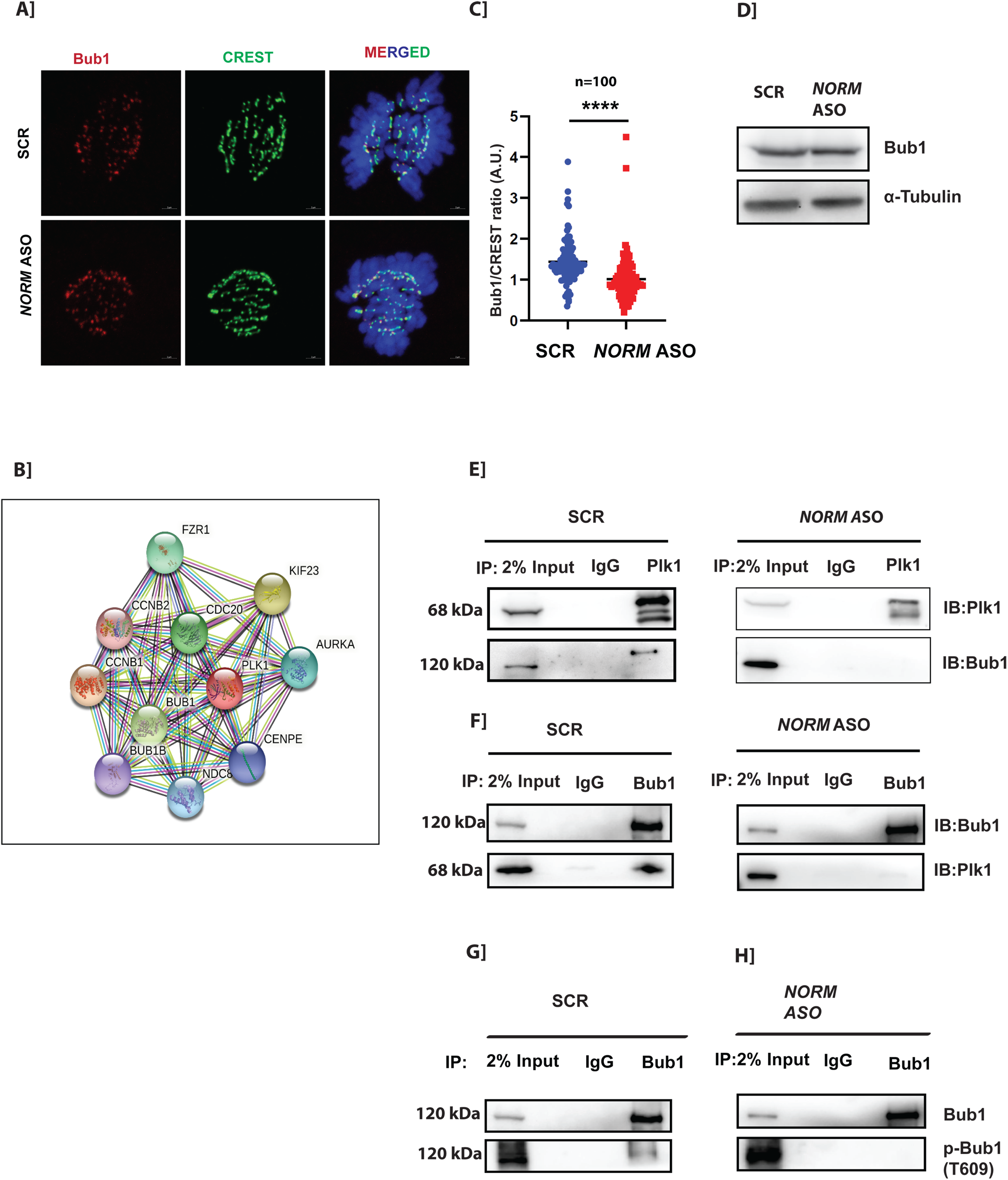

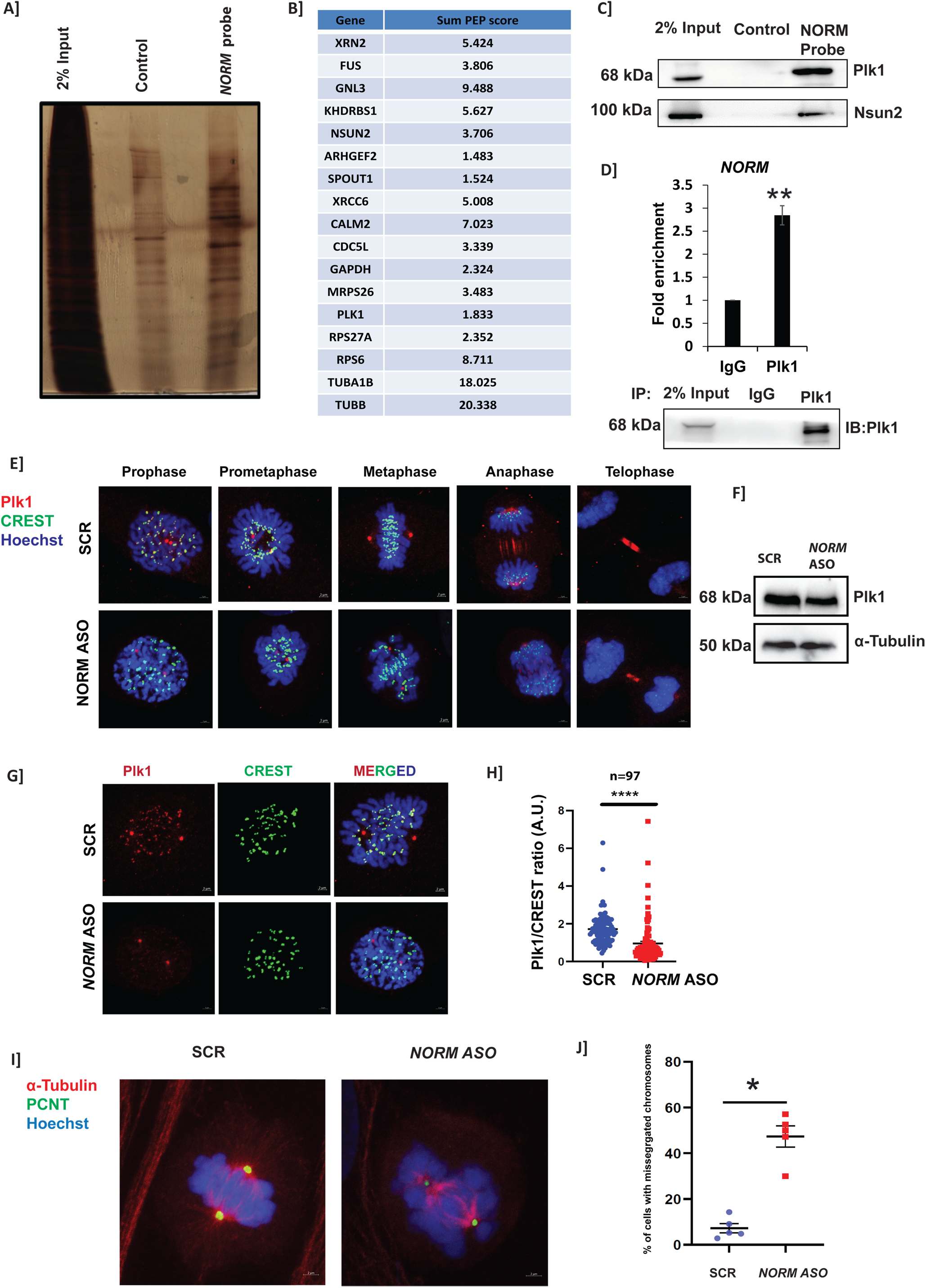

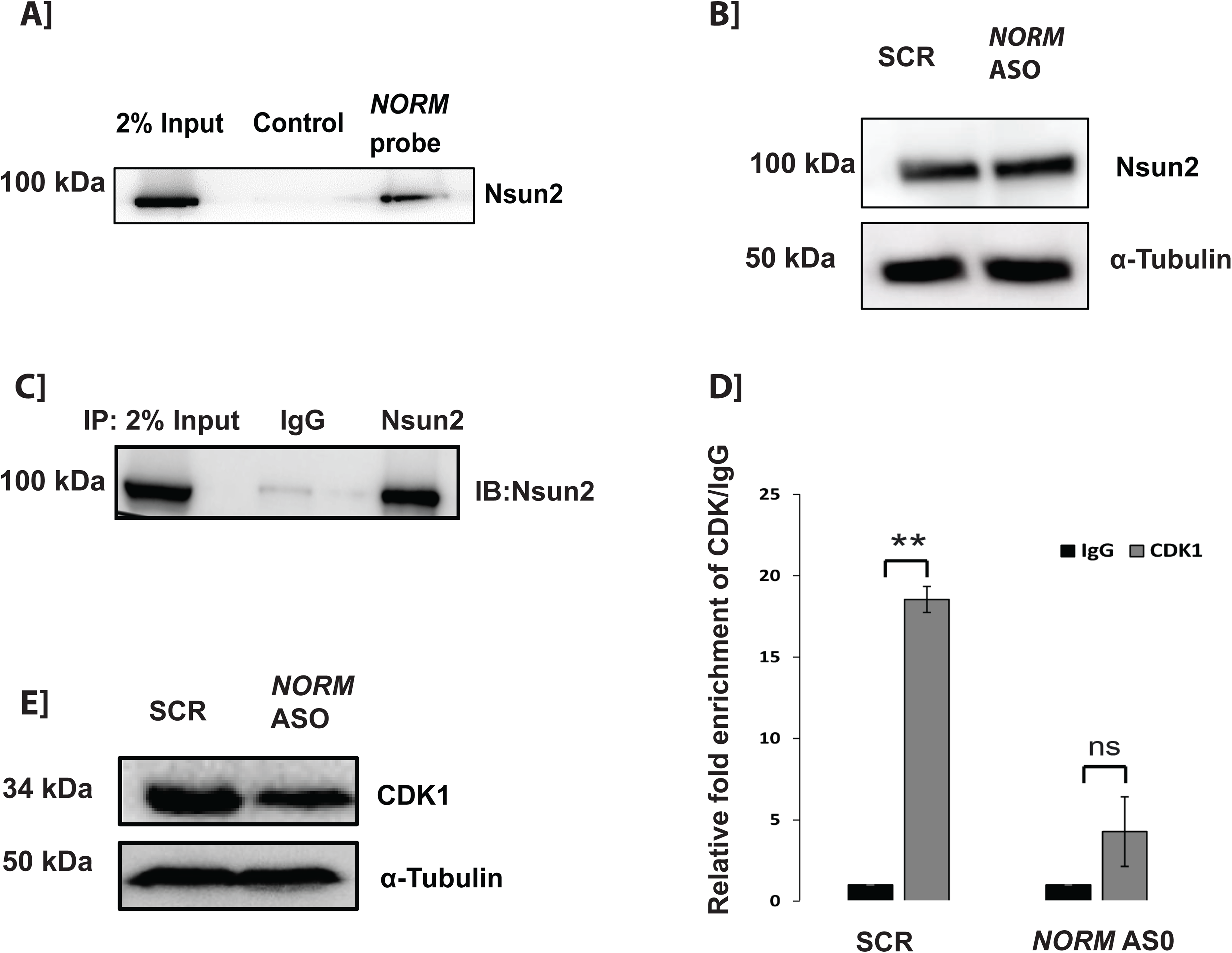

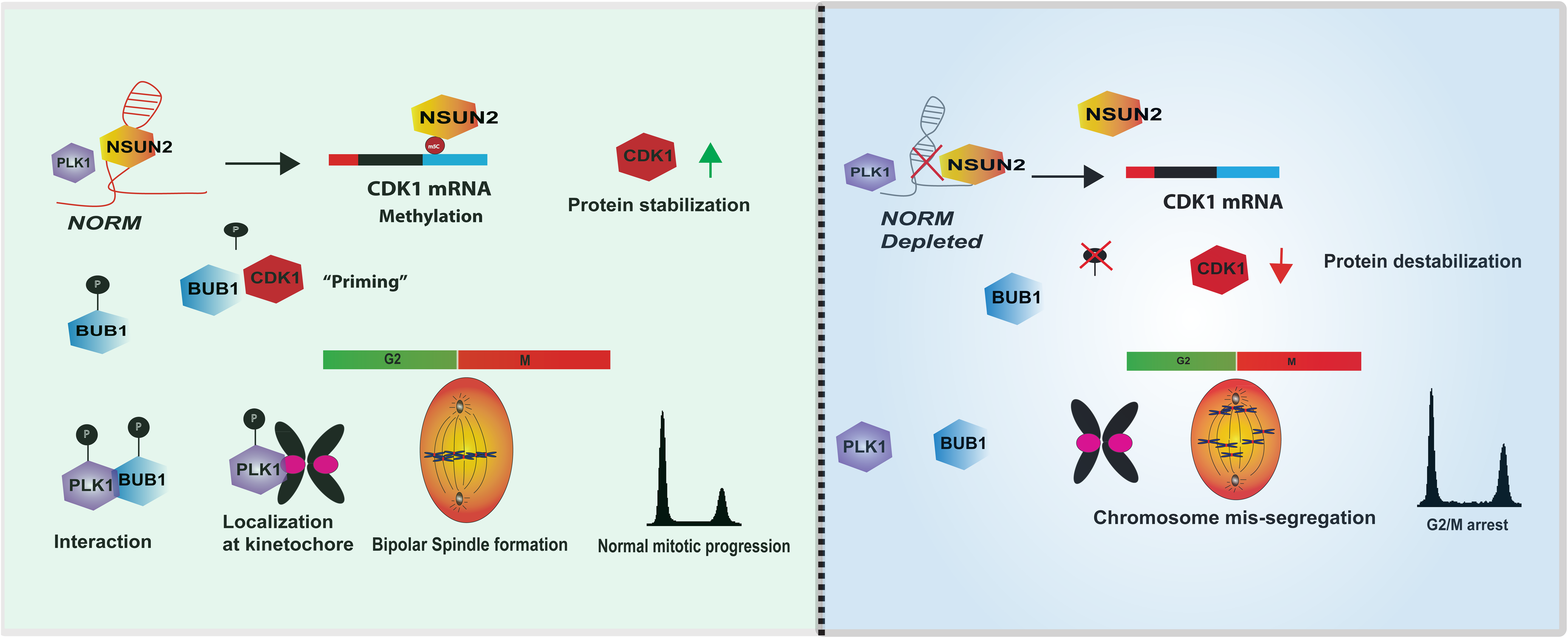

