## supplementary data 1 for "lncRNA *NORM* is essential for proper chromosome segregation through the Plk1-Bub1 and Nsun2 axis"

A]

| Gene | Log2(Fold change ) |
|---|---|
| SEC31A | 10.18208 |
| ITGA2 | 9.846807 |
| FBXO6 | 6.806259 |
| DNM2 | 6.543368 |
| SMURF1 | 6.512652 |
| GUCD1 | 5.74763 |
| WDR4 | 5.743208 |
| POP1 | 5.738942 |
| PPP1R7 | 5.604307 |
| PSMD1 | 5.515586 |

| Gene | Log2(Fold change ) |
|---|---|
| ZNF48 | -9.10621 |
| SLC16A6 | -8.57854 |
| BRCA1 | -8.18513 |
| CHRND | -7.22453 |
| PCNT | -7.15993 |
| DCAKD | -7.06604 |
| POT1 | -6.9355 |
| ZNF74 | -6.67736 |
| RNF14 | -6.64547 |
| YY1AP1 | -6.39752 |

B]

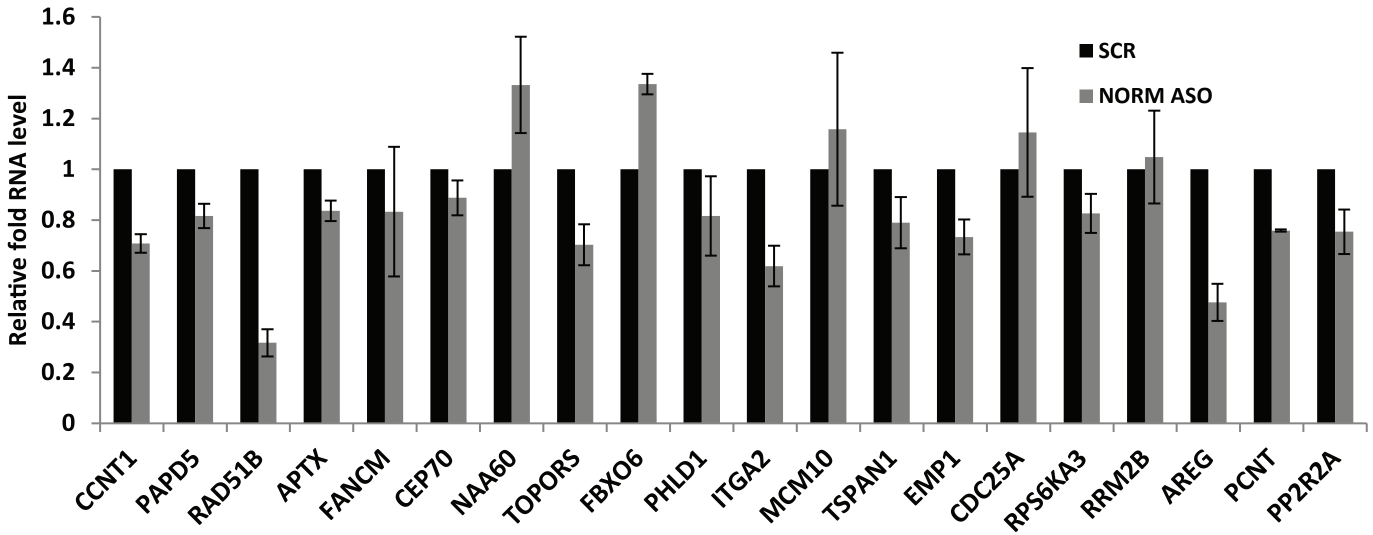
