## supplementary data 2 for "lncRNA *NORM* is essential for proper chromosome segregation through the Plk1-Bub1 and Nsun2 axis"

qPCR primer List

| **Gene** | **Sequence (5’🡪3’)** |
| --- | --- |
| **qPCR primers** | |
| HPRT FP | GAAAAGGACCCCACGAAGTGT |
| HPRT RP | AGTCAAGGGCATATCCTACAACA |
| *NORM* FP | CTCACCAACCGTGGAAATAA |
| *NORM* RP | GGCCAGTCTCCTTCTGTGATG |
| RNR1 FP | ATATACCGCCATCTTCAGCAAAC |
| RNR1 RP | CCCATTTCTTGCCACCTCAT |
| MALAT1 FP | TTCCGGGTGTTGTAGGTTTC |
| MALAT1 RP | TCTCCAGGACTTGGCAGTCT |
| CCNT1 FP | ACAACAAACGGTGGTATTTCACT |
| CCNT1 RP | CCTGCTGGCGATAAGAAAGTT |
| APTX FP | CAAAGGCCCGTTACCATTGG |
| APTX RP | ACCCAGCAAAATCTACAATCACC |
| FANCM FP | CAACAACTCCACCGGATAGTG |
| FANCM RP | GTTGGCCTCCTTACTGCAATC |
| TSPAN1 FP | AATCTTGGCTCTAAGTGCCAC |
| TSPAN1 RP | TCTGCCCAATTAGCAGGTTAGTA |
| KRT16 FP | GACCGGCGGAGATGTGAAC |
| KRT16 RP | CTGCTCGTACTGGTCACGC |
| NAA60 FP | CCAACTTCTCTGTTGACACACA |
| NAA60 RP | AGTAAGAGGGAACCTATGCCG |
| ITGA2 FP | CCTACAATGTTGGTCTCCCAGA |
| ITGA2 RP | AGTAACCAGTTGCCTTTTGGATT |
| AREG FP | GAGCCGACTATGACTACTCAGA |
| AREG RP | TCACTTTCCGTCTTGTTTTGGG |
| EMP1 FP | GTGCTGGCTGTGCATTCTTG |
| EMP1 RP | CCGTGGTGATACTGCGTTCC |
| PHLDA1 FP | GGAGATCGACTTTCGGTGCC |
| PHLDA1 RP | GGCCTGACGATTCTTGTACTG |
| PAPD5 FP | GACATCGACCTAGTGGTGTTTG |
| PAPD5 RP | CGACTTTGTGTTTCCGAAGAGC |
| CDC25A FP | GTGAAGGCGCTATTTGGCG |
| CDC25A RP | TGGTTGCTCATAATCACTGCC |
| RPS6KA3 FP | AGGGCAGGGATCATTTGGAAA |
| RPS6KA3 RP | GGCCTTCTTCAATACCTTCATGG |
| RAD51B FP | CAGAGTCTGCATTTAGTGCTGA |
| RAD51B RP | ACAGGTGAGTTCCCGATAAAGAT |
| RRM2B FP | ATTGGGCCTTGCGATGGATAG |
| RRM2B RP | GAGTCCTGGCATAAGACCTCT |
| MCM10 FP | CCCCTACAGACGATTTCTCGG |
| MCM10 RP | CAGATGGGTTGAGTCGTTTCC |
| PPP2R2A FP | GCGAGACATAACCCTAGAAGC |
| PPP2R2A RP | CACTTGCACAGACTTTGCGAG |
| CEP70 FP | AGAACAACGAGCTAATGACTTGG |
| CEP70 RP | AGCCCTACTTAGTGATTCATCCT |
| PCNT FP | AGGAGGAGAGTCCGGTAACC |
| PCNT RP | TCAGGGGTGTCGTCACATGAT |
